# High-throughput single-cell transcriptome profiling of plant cell types

**DOI:** 10.1101/402966

**Authors:** Christine N. Shulse, Benjamin J. Cole, Gina M. Turco, Yiwen Zhu, Siobhan M. Brady, Diane E. Dickel

**Affiliations:** Department of Energy Joint Genome Institute, Walnut Creek, CA, 94598 USA.; Department of Plant Biology and Genome Center, UC Davis, Davis, CA 95616 USA.; Environmental Genomics and Systems Biology Division, Lawrence Berkeley National Laboratory, Berkeley, CA 94720, USA.

## Abstract

Single-cell transcriptome analysis of heterogeneous tissues can provide high-resolution windows into the genomic basis and spatiotemporal dynamics of developmental processes. Here we demonstrate the feasibility of high-throughput single-cell RNA sequencing of plant tissue using the Drop-seq approach. Profiling of >4,000 individual cells from the *Arabidopsis* root provides transcriptomes and marker genes for a diversity of cell types and illuminates the gene expression changes that occur across endodermis development.

---

Single-cell transcriptomics technologies are revolutionizing molecular studies of heterogeneous tissues and organs, enabling the elucidation of new cell type populations and revealing the cellular underpinnings of key developmental processes^1–3^. Recently developed high-throughput single-cell RNA-seq (scRNA-seq) techniques, such as Drop-seq^4^, use a microfluidic device to encapsulate cells in emulsified droplets, allowing for the profiling of hundreds or even thousands of cells in a single experiment. Despite this remarkable advance, the large and non-uniform size of plant cells, as well as the presence of cell walls, have hindered the application of this technology to plant tissues. Applying high-throughput scRNA-seq methods to plants would negate the need for specialized reporter lines that are currently widely used for the capture of specific cell type populations. Single-cell technologies have the potential to provide a detailed spatiotemporal characterization of distinct cell types present in plants, along with the transcriptional pathways that regulate their diversity and development^5^.

As a proof of principle for high-throughput, microfluidics-enabled scRNA-seq in plant tissue, we performed Drop-seq on protoplasts isolated from 5-day-old *Arabidopsis thaliana* roots (**Fig. 1** and **Supplementary Table 1**). Across two technical replicates, we obtained transcriptomes for a total of 4,043 individual root cells, with a minimum of 1,000 unique molecular identifier (UMI)-tagged transcripts per cell (**Supplementary Figure 1a,b** and **Methods**). To confirm that Drop-seq captured a representative population of cells present in the root, we combined the expression of all captured cells into a pseudo-bulk profile and compared this profile to a conventional mRNA-seq profile of non-protoplasted 5-day old *Arabidopsis* root tissue (**Supplementary Fig. 1c,d**). The pseudo-bulk transcriptome showed high correlation with the bulk root mRNA-seq profile (Spearman’s rho: 0.74 for all genes, 0.75 when known protoplast response genes^6^ were excluded) and much lower correlation with a previously reported^7^ bulk whole flower mRNA-seq profile (Spearman’s rho: 0.38-0.40).

**Figure 1.**
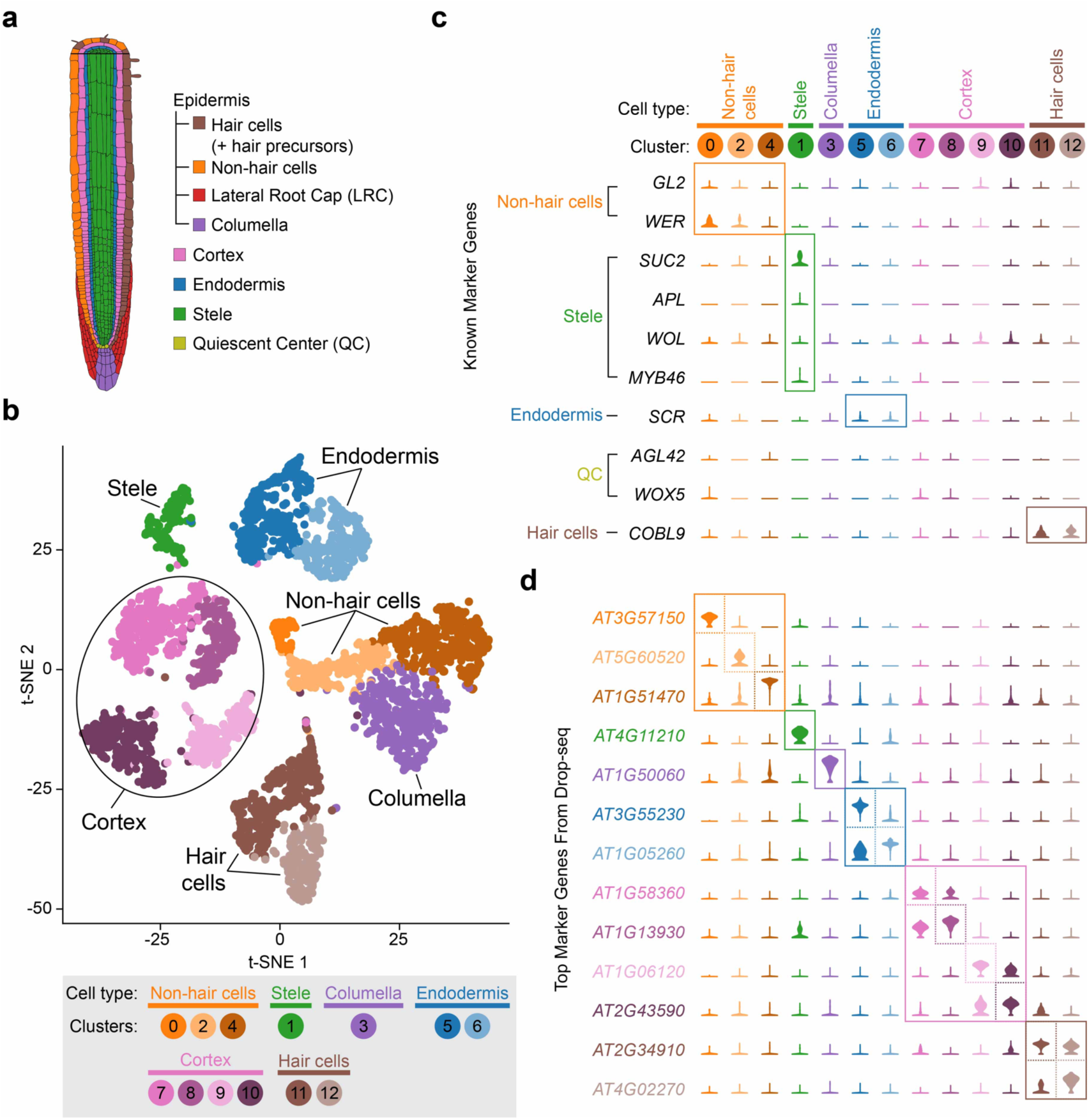
Single-cell RNA-seq of 4,043 *Arabidopsis* root cells captures diverse cell types. **a)** Cartoon representing the cell types that comprise the *Arabidopsis* root. **b)** t-SNE dimensional reduction of 4,043 single Arabidopsis root cells that were profiled using Drop-seq. Cells were clustered into 13 populations using Seurat8. Points indicate individual cells and are colored by assigned cell type and cluster according to the legend below (see also **Methods** and **Supplementary Figs. 2-3** for details of how clustering was performed and cell type identities assigned). **c)** Violin plots showing the expression of ten previously identified cell type marker genes across all clusters. Labels to the left of the gene names indicate the cell population each gene is used to designate. **d)** Violin plots showing the cluster-specific expression of the top-ranking candidate novel marker gene for each cell population identified by this study, color-coded as in **b** and **c**.

To identify distinct cell populations, we performed unsupervised clustering analysis of the 4,043 single cells with Seurat^8^ (**Fig. 1b** and **Supplementary Fig. 2a**). All 13 clusters identified contained cells from each technical replicate in similar proportions, supporting high reproducibility of the approach and an absence of substantial batch effects in our experiments (**Supplementary Fig. 2b,c**). To assign individual clusters to known major root cell types, we examined the expression of commonly used marker genes^9^ in our single-cell profiles and computed an Index of Cell Identity (ICI) score^10^ for each cell (**Fig. 1b,c** and **Supplementary Fig. 3a-c**; see also **Methods**). Nearly all major cell types present in the root were captured by at least one cluster, including stele (one cluster), endodermis (two clusters), cortex (four clusters), hair cells (two clusters), non-hair cells (three clusters), and columella (one cluster). We could not assign a distinct cell cluster to quiescent center (QC) cells, likely due to their rarity^11^. Additionally, we did not assign a cluster to lateral root cap cells, as this cell type is not included in the ICI model and because of the paucity of known marker genes that specifically define this cell type. We compared the proportions of each cell type captured by Drop-seq to previous microscopy-based cell population surveys of root cell types across developmental time^11^. This highlighted a relative depletion of stele cells in our data, possibly due to inefficient dissociation or cell death of this population during protoplasting (**Supplementary Fig. 3d**). Aside from stele, other cell types were well captured by Drop-seq. Overall these results demonstrate that Drop-seq accurately captures the complexity of cell types present in the plant root.

Expression analysis of marker genes commonly used to label specific cell populations revealed many examples of genes with remarkably limited specificity or very low expression levels (**Fig. 1c**). Therefore, we analyzed the scRNA-seq data to identify a new set of robust marker genes, quantitatively optimized for specificity. The top candidate marker gene for each cluster is shown in **Fig. 1d**, with the top 50 candidate marker genes for each cluster listed in **Supplementary Table 2**. This unbiased approach identified genes with previously established roles in the biology of some of the cell populations. For example, all five genes encoding components of the Casparian strip (*CASP1* through *CASP5*) are among the top 20 marker genes for one of the endodermis clusters (Cluster 5), and multiple root hair-specific genes (e.g., *COBL9, EXPA7, EXPA18*, and *ADF8*) are top candidate marker genes for one or both hair cell clusters (Clusters 11 and 12). However, genes with well-established roles in root development represent a relatively small subset of the highly specific marker genes we identified, and many of the candidate marker genes have little to no functional data presently available. As evidence of this, 43% of the top 50 marker genes (**Supplementary Table 2**), including 7 of the 13 top (rank 1) marker genes across all clusters, have no gene symbol beyond an AGI Gene Identifier. Taken together, these marker genes provide a number of new avenues to explore the transcriptional processes that underlie root cell type specification and function.

A particular strength of the application of scRNA-seq to developing tissues is the ability to infer developmental trajectories of individual cell populations through pseudotime analysis^13^. Individual roots form an apical-basal anatomical gradient of temporal differentiation states, with undifferentiated (i.e., meristematic) cells near the root tip and mature cells located nearer the root-shoot junction. Therefore, we hypothesized that the various subpopulations identified for some cell types may reflect these trajectories. We carried out a pseudotime analysis, focusing first on endodermal cells, which have a known trajectory from undifferentiated, to state I (defined by the formation the Casparian strip), to state II (defined by the formation of a secondary cell wall made of suberin)^14^ (**Fig. 2a**). In this analysis, we isolated and re-embedded the scRNA-seq profiles of all 698 endodermis cells belonging to clusters 5 and 6 into a new t-SNE, then defined a pseudotime axis between the centroids of the two clusters (**Fig. 2a,b**, see also **Methods**). In order to properly orient the pseudotime axis relative to known endodermis differentiation, we 1) examined the expression profiles of genes known to be dynamically expressed during endodermal development (**Fig. 2c** and **Supplementary Fig. 4a**); and 2) compared our pseudotime expression profiles to early and mature endodermis expression data generated previously from marker gene lines^15^ (**Supplementary Fig. 4b,c**). Once we established the correctly oriented pseudotime trajectory, we used it to ascertain additional genes that are dynamically regulated during endodermis development. Searching for genes with a high expression variance (variance > 0.2) over pseudotime identified 270 total dynamically expressed genes, which fell into three patterns: early, middle, and late expression (**Fig. 2d** and **Supplementary Table 3**). The early expression gene group includes *MYB36*, which is known to regulate the initial stages of endodermis differentiation^16^, and this group is enriched for gene ontology (GO) terms associated with water or fluid transport (**Fig. 2d** and **Supplementary Table 4**). Genes whose expression peaks in the middle of the pseudotime axis include canonical state I genes (e.g., *CASPs, ESB1*) and are enriched for GO terms associated with Casparian strip assembly. Genes whose expression peaks late are enriched in GO terms associated with response to chemicals and stress, consistent with the role of stress hormones in the induction of state II differentiation^17^. Currently, the specific transcriptional pathways that directly regulate state II differentiation are not well defined^14^. Thus, these late expressing genes are candidates for the cellular components that direct mature endodermis differentiation. Transcription factor genes found in this late expressing group include *WRKY33, TBF1, ABR1, ATAF2,* and *AT5G01380*. Additionally, numerous fundamental questions in endodermis biology remain outstanding, including the identification of genes that regulate or contribute to the biosynthesis of lignin and suberin^18^. Intriguingly, 54 of the genes with dynamic expression across endodermis development have predicted oxidoreductase activity, a molecular function implicated in the polymerization of these compounds^19^ (**Supplementary Table 3**). We performed a similar pseudotime analysis for hair cells (clusters 11 and 12) and identified 400 developmentally dynamic genes, including many with roles in actin and cytoskeleton reorganization (**Supplementary Fig. 5** and **Supplementary Tables 5-6**). Collectively, these results highlight the power of scRNA-seq for generating new hypotheses that may lead to a deeper understanding of cellular development in plant tissues.

**Figure 2.**
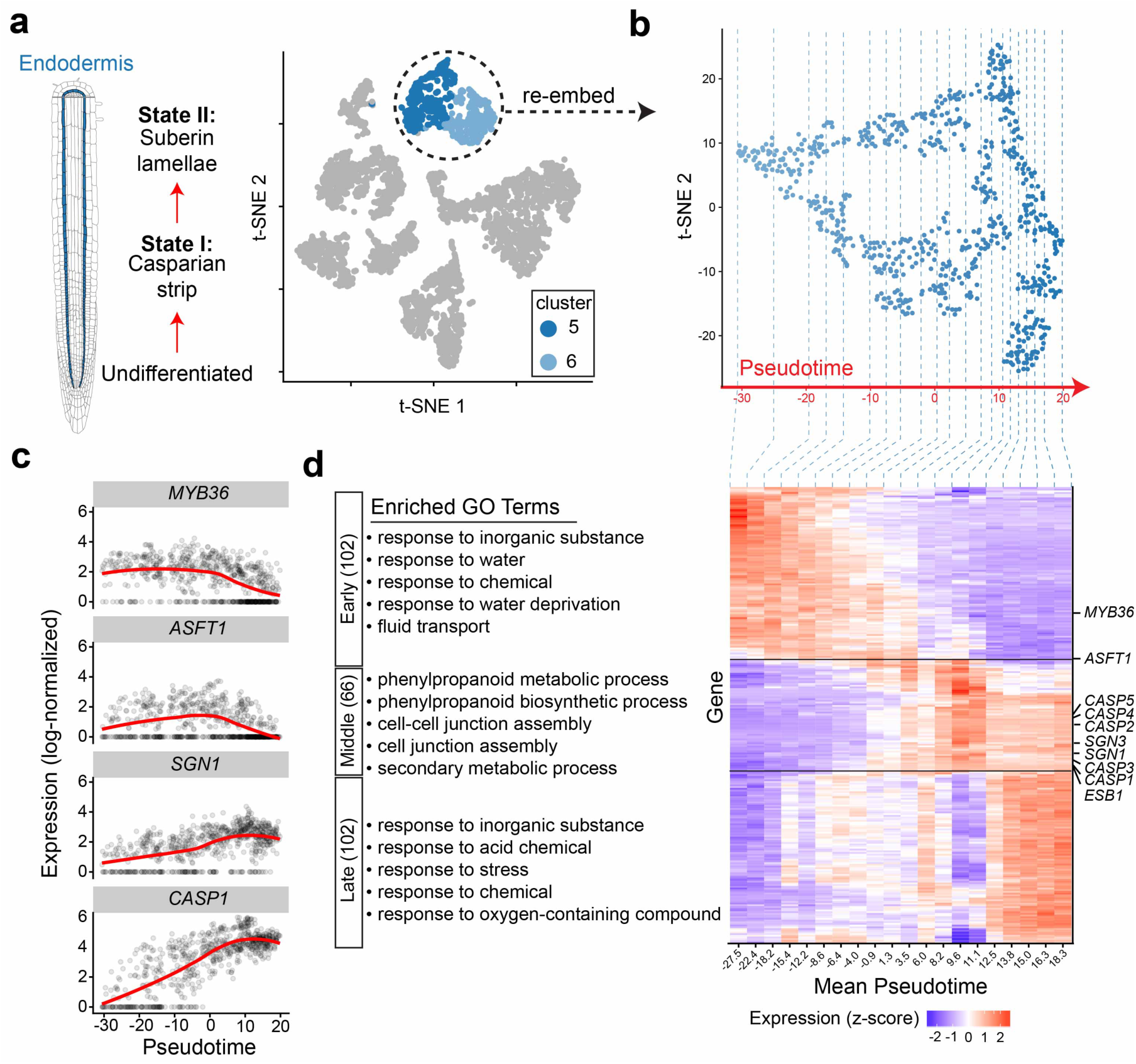
Transcriptional dynamics of endodermis development. **a)** Left: Schematic representation of the trajectory of endodermis development. **Right:** t-SNE representation, as in **Fig. 1b**, with only the endodermis cells colored blue. **b)** The 698 endodermis cells were embedded in a new t-SNE dimensional space, and a pseudotime axis was derived empirically (see **Methods**). Each point indicates a unique cell. **c)** Expression profiles along pseudotime for genes known to have dynamic expression during endodermal development. Each point indicates the log-normalized expression of the indicated gene in a single cell, with the LOESS regression line over pseudotime in red. **d)** Right: Heat map showing the gene expression z-scores of 270 developmentally regulated endodermis genes identified by scRNA-seq (see **Methods**). For this plot, all endodermis cells were divided into 20 bins of 34-35 cells each. Canonical endodermal development genes are noted along the right axis. Genes were grouped by expression pattern, with groups delineated by horizontal black lines. From top to bottom: early, middle, and late expression). Left: top five gene ontology (GO) terms enriched in each gene expression group. Numbers in parentheses indicate the number of genes in each category.

This study demonstrates that Drop-seq is a powerful method to quickly profile the diverse cell types that comprise heterogenous plant tissue. In just two experiments, we obtained scRNA-seq profiles for a few thousand cells, exceeding the throughput of previously reported fluorescence-activated cell sorting (FACS)-based scRNA-seq by more than an order of magnitude^3^. Our results suggest that Drop-seq will be applicable to any tissue or plant species for which protoplasts can be obtained. We anticipate that the dataset described herein will be a valuable resource to other investigators for the study of root cell type differentiation and that Drop-seq will open exciting new inroads of exploration into the identities and functions of plant cell types.

## Methods

### Plant material and growth conditions

Arabidopsis (*Arabidopsis thaliana*) ecotype Columbia (Col-0) seeds were surface sterilized in 70% ethanol for 2 minutes, followed by 50% bleach and 0.1% Triton X-100 for 5 - 10 minutes. Sterilized seeds were washed 4 - 6 times with sterile water and imbibed in the dark for 3 days at 4°C. Seeds were sown at a density of approximately 125 seeds per row on sterile nylon mesh filters placed on top of plant growth media consisting of 1X Murashige and Skoog Basal Medium (Sigma-Aldrich, St. Louis, MO) supplemented with 1% sucrose (Sigma-Aldrich), 1% agar (Sigma-Aldrich), 2.6 mM 2-(N-Morpholino)ethanesulfonic acid (MES; Sigma-Aldrich), and adjusted to pH 5.7. Petri dishes were positioned vertically in an incubator (Percival Scientific, Perry, IA) under a long-day photoperiod (16 h of light) at 22°C.

### Drop-seq of *Arabidopsis* root protoplasts

Protoplast suspensions for the two Drop-seq runs described here were prepared from approximately 10,000 5-day-old *Arabidopsis* seedlings using previously described methods^20^. Briefly, *Arabidopsis* roots were finely chopped and distributed among seven 70-μm cell strainers (each containing 1,250-1,500 roots). Each strainer was then immersed in a solution of cellulysin (Sigma-Aldrich) and pectolyase (Sigma-Aldrich) for 60 minutes. After cell wall digestion, each strainer, containing remnants of roots that were not protoplasted, was removed and discarded. Protoplasts were then pelleted, resuspended, and filtered through a 40-μm cell strainer to eliminate any clumped cells. The cells were then kept either at their starting concentration (137 cells/μl) or diluted in Solution A^20^ (10 mM KCl, 2 mM MgCl_2_, 2 mM CaCl_2_, 0.1% bovine serum albumin, 2 mM MES, 600 mM mannitol) to 97 cells/μl. Human cells were spiked in at a concentration of 2.5%, and Drop-seq was performed as described previously in Reference 4 and Drop-seq Laboratory Protocol version 3.1 (http://mccarrolllab.com/dropseq/). The two libraries were uniquely indexed and then pooled and sequenced on a single lane of a HiSeq 4000.

### Read alignment and generation of digital gene expression data

A digital expression matrix of Single-cell Transcriptomes Attached to MicroParticles (STAMPs) was created from the raw sequencing data as described in the Drop-seq Core Computational Protocol version 1.0.1 (http://mccarrolllab.com/dropseq/). Sequence data was aligned to a combined human (GRCh38)-*Arabidopsis thaliana* (TAIR 10) mega-reference using STAR v2.5.2b with the default settings. Uniquely mapped reads were grouped by cell barcode. To identify true STAMPs, we discarded any cell barcodes with fewer than 1,000 unique transcripts (i.e., UMIs). A STAMP was then considered as derived from *Arabidopsis* rather than human or mixed if 98% or more of the mapped reads mapped to *Arabidopsis*.

### Principal component and clustering analysis

Data from two independent Drop-seq runs (using the 137 and 97 cells/µL preparations of the same protoplast prep of 5-day old Arabidopsis roots, respectively) were combined into a single data set for further analysis. After removing genes known to be induced during protoplasting^6^, the Seurat R package^8^ (version 2.3.1) was used to log-normalize, scale, and identify highly variable genes (1,002 in total) among the scRNA expression profiles. Seurat was then used to compute 30 principal components (PCs). Informative PCs were inferred based on the JackStraw analysis function in the Seurat package. All PCs up to but not including the first PC having a p-value less than 0.01 were defined as informative. The informative PCs (30 in the case of the combined 5-day root scRNA-seq data) were used to compute a t-SNE^21^ dimension reduction, and to group cells into 13 distinct clusters. Clustering was performed with the FindClusters function in the Seurat package, using a resolution of 0.6 and default parameters. For each cluster, the average expression of the 1,002 highly variable genes was calculated over all member cells. These expression profiles were then used to re-order the numerical cluster assignments based on hierarchical clustering and leaf order optimization using the Seriation^22^ package (version 1.2-3). For each cluster of cells, new putative marker genes were computed using the FindAllMarkers function in Seurat.

### Assignment of clusters to known cell types

To align cell population clusters from the unsupervised scRNA-seq to known cell types, we assessed 1) expression of known cell-type specific marker genes and 2) Index of Cell Identity (ICI) scores^10^. For marker gene expression-guided assignments, a set of known or inferred marker genes was identified from the literature (e.g., Ref. 7). The average normalized expression for these genes for each cell was computed and used to guide assignments of cell clusters to putative cell types. We next used ICI, which assesses the expression of hundreds of genes to calculate a probability that a single cell belongs to specific *Arabidopsis* root cell type and returns the best cell type match. Probabilities associated with the cell type assignments were estimated using a bootstrapped permutation method (1000 iterations) and adjusted using the Benjamini-Hochberg method^23^. Clusters of cells within the t-SNE graph were assigned an overall cell type based on the preponderance of ICI calls, regardless of their significance (**Supplementary Fig. 3a,b**), however more weight was given to cells with significant adjusted p-values (p < 0.01, **Supplementary Fig. 3c**).

### Pseudotime analysis

To characterize the developmental trajectory of endodermis, a “pseudotime” axis^24^ (developmental state along a progression) was derived for these cells. Due to the low-complexity nature of the developmental states predicted in the endodermis, we opted to compute a simplistic developmental trajectory. To do this, we first isolated the scRNA-seq profiles of all endodermis cells (clusters 5 and 6) and re-computed the PCA and t-SNE dimension reductions for each reduced set. Next, we defined the “pseudotime” axis as the straight line between the centroid of cluster 5 and the centroid of cluster 6. As t-SNE axes are arbitrary, we then rotated the endodermis t-SNE such that the x-axis corresponded to this pseudotime axis. All cells within clusters 5 and 6 were then assigned a pseudotime value based on their position along the pseudotime line. Endodermis pseudotime was then divided into 20 bins, such that each bin had approximately the same number of cells (34-35 each). The log-normalized expression, as well as the z-score (scaled and centered gene expression) was then computed for each gene among these bins. Genes with a normalized expression variance above 0.2 were considered developmentally regulated. An identical procedure was used to characterize the developmental trajectory of hair cells (clusters 11 and 12).

### Analysis of enriched GO terms

A PANTHER overrepresentation test^12^ was performed using a list of genes identified for each expression pattern (endodermis: early, middle, and late; trichoblasts: early and late) against the *Arabidopsis thaliana* genome.

### Bulk root tissue mRNA-seq and Pseudo-bulk analysis

To compare Drop-seq expression data to standard mRNA-seq data, we isolated total RNA from flash frozen 5-day-old Arabidopsis root material using the Qiagen RNeasy kit (Carlsbad, CA). The RNA was used in the TruSeq Stranded mRNA Library Prep Kit (Illumina, RS-122-2101) according to the manufacturer’s Low Sample (LS) Protocol. The library was uniquely indexed and then sequenced on a single lane of a HiSeq 4000. Transcript counts for each gene were then tabulated. In parallel, a pseudo-bulk data set was generated from the Digital Gene Expression matrix from scRNA-seq of the root protoplasts. A separate transcript count profile from Arabidopsis floral bud tissue was also downloaded from the NCBI GEO database (GSM2616967). These three expression profiles (filtered to contain only genes with FPKM > 1) were then normalized together using the DESeq2 package^25^. The Spearman correlation coefficients between the pseudo-bulk and whole-root and whole-flower data sets were computed in R before or after filtering out known protoplast-inducible genes.

### Data availability

Data have been deposited in NCBI’s Gene Expression Omnibus: accession number GSE116614.

## Acknowledgements

This work was supported by a grant to D.E.D. from the Laboratory Directed Research and Development Program of Lawrence Berkeley National Laboratory. Work was performed at LBNL and the US DOE Joint Genome Institute under U.S. Department of Energy Contract No. DE-AC02-05CH11231. G.M.T. and S.M.B. were funded by a Howard Hughes Medical Institute Faculty Scholar Fellowship. G.M.T. was additionally funded by an NSF predoctoral fellowship and the AAUW Dissertation Completion Fellowship. This work used the Vincent J. Coates Genomics Sequencing Laboratory at UC Berkeley, supported by NIH S10 OD018174 Instrumentation Grant. We thank V. Singan for generating library counts from raw mRNA-Seq data and I. Barozzi for computational advice. The authors would like to acknowledge E. Macosko, M. Goldman, J. Nemesh, and S. McCarroll for providing detailed Drop-seq protocols. Additionally, thank you to N. Geldner for discussing endodermis pseudotime ordering and to A. Visel, R. O’Malley, and J. Vogel for helpful feedback on the manuscript.

## Author Contributions

C.N.S. and Y.Z. performed experiments. C.N.S. and B.J.C. carried out data analysis. G.M.T. and S.M.B. contributed expert guidance on protoplasting and plant root development. D.E.D. and S.M.B. oversaw the study. All authors contributed to the preparation of the manuscript.

